# The Detection of Phase Amplitude Coupling During Sensory Processing

**DOI:** 10.1101/163006

**Authors:** R.A Seymour, G. Rippon, K. Kessler

## Abstract

There is increasing interest in understanding how the phase and amplitude of distinct neural oscillations might interact to support dynamic communication within the brain. In particular, previous work has demonstrated a coupling between the phase of low frequency oscillations and the amplitude (or power) of high frequency oscillations during certain tasks, termed phase amplitude coupling (PAC). For instance, during visual processing in humans, PAC has been reliably observed between ongoing alpha (8-13Hz) and gamma-band (>40Hz) activity. However, the application of PAC metrics to electrophysiological data can be challenging due to numerous methodological issues and lack of coherent approaches within the field. Therefore, in this article we outline the various analysis steps involved in detecting PAC, using an openly available MEG dataset from 16 participants performing an interactive visual task. Firstly, we localised gamma and alpha-band power using the Fieldtrip toolbox, and extracted time courses from area V1, defined using a multimodal parcellation scheme. These V1 responses were analysed for changes in alpha-gamma PAC, using four common algorithms. Results showed an increase in gamma (40-100Hz) - alpha (7-13Hz) PAC in response to the visual grating stimulus, though specific patterns of coupling were somewhat dependent upon the algorithm employed. Additionally, post-hoc analyses showed that these results were not driven by the presence of non-sinusoidal oscillations, and that trial length was sufficient to obtain reliable PAC estimates. Finally, throughout the article, methodological issues and practical guidelines for ongoing PAC research will be discussed.

## 2. Introduction

Electrophysiological brain oscillations are often separated into distinct frequency bands, ranging from low-frequency delta (1-4Hz) to high-frequency gamma (<40Hz). The power and/or connectivity profiles of these frequency bands have been linked with specific neuronal and cognitive functions (Buzsáki & Draguhn, 2004; Palva, Palva, & Kaila, 2005). Whilst this has proven a powerful tool in neuroscientific research, there is emerging evidence that oscillations from different frequency bands also display specific coupling patterns – a phenomenon termed cross frequency coupling (CFC) (Hyafil, Giraud, Fontolan, & Gutkin, 2015; Jensen & Colgin, 2007). One of the best studied forms of CFC is phase-amplitude coupling (PAC), in which the amplitude/power of a high frequency oscillation, often gamma (>40Hz), is coupled to the phase of a lower frequency oscillation (Canolty et al., 2006; Canolty & Knight, 2010). PAC has been observed in multiple regions of the human brain, including the visual cortex (Voytek et al., 2010), auditory cortex (Cho et al., 2015), hippocampus (Heusser, Poeppel, Ezzyat, & Davachi, 2016; Lega, Burke, Jacobs, & Kahana, 2014) and prefrontal cortex (Voloh, Valiante, Everling, & Womelsdorf, 2015; Voytek et al., 2015), in both electrocorticography (ECOG) and magnetoencephalography (MEG) recordings.

Within the visual system, there is strong evidence for a dynamic coupling between alpha phase (8-13Hz) and gamma amplitude (>40Hz) (Bonnefond & Jensen, 2015; Spaak, Bonnefond, Maier, Leopold, & Jensen, 2012; Voytek et al., 2010). Alpha oscillations are associated with pulses of cortical inhibition every ∼100ms (Jensen & Mazaheri, 2010; Klimesch, 2012), whilst supporting communication through phase dynamics (Fries, 2015). In contrast, gamma oscillations emerge through local excitatory and inhibitory interactions, and synchronise local patterns of cortical activity (Buzsáki & Wang, 2012; Singer & Gray, 1995). In visual cortex, ongoing gamma-band activity becomes temporally segmented by distinct phases of alpha-band activity (Bonnefond, Kastner, & Jensen, 2017; Spaak et al., 2012), possibly via inter-laminar coupling between supragranular and infragranular cortical layers (Mejias, Murray, Kennedy, & Wang, 2016). Intriguingly, this coupling has been proposed to act as a mechanism for the dynamic coordination of brain activity over multiple spatial scales, with high-frequency activity within local ensembles coupled to large-scale patterns of low-frequency phase synchrony (Bonnefond et al., 2017), both within the visual system (Bonnefond & Jensen, 2015), and more widespread neurocognitive networks (Florin & Baillet, 2015). This would allow information to be routed efficiently between areas and for neuronal representations to be segmented and maintained, for example during visual working memory (Bonnefond & Jensen, 2015; Lisman & Idiart, 1995). Atypical patterns of PAC have also been proposed to underlie atypical cortical connectivity in several neurological conditions, including autism spectrum disorder (Kessler, Seymour, & Rippon, 2016; Khan et al., 2013), schizophrenia (Kirihara, Rissling, Swerdlow, Braff, & Light, 2012) and Parkinson’s Disease (De Hemptinne et al., 2013; Özkurt & Schnitzler, 2011).

Given the developing interest in cross-frequency coupling, it is vital for the wider neuroscience and electrophysiological community to understand the steps involved in its measurement and interpretation. This is especially important for PAC, which is beset with methodological pitfalls, since there are many competing algorithms, approaches, and currently no gold-standard set of analysis steps (Canolty & Knight, 2010; Jensen, Spaak, & Park, 2016). It has also been suggested that numerous incidences of reported PAC may in fact be false positives, caused by suboptimal analysis practices and/or the presence of artefacts within the data (Aru et al., 2015; Hyafil, 2015). For example non-sinusoidal sawtooth-like oscillations can generate artificially inflated PAC values, via low-frequency phase harmonics (Cole et al., 2017; Lozano-Soldevilla, ter Huurne, & Oostenveld, 2016; Vaz, Yaffe, Wittig Jr, Inati, & Zaghloul, 2017).

In this article, we outline a general approach for detecting changes in phase-amplitude coupling during visual processing, using a novel MEG dataset, analysed using the Fieldtrip toolbox (Oostenveld, Fries, Maris, & Schoffelen, 2010), and openly available MATLAB scripts. Four common PAC algorithms were used to quantify the coupling between ongoing alpha phase (7-13Hz) and gamma amplitude/power (>40Hz) whilst participants viewed a static visual grating. Given the controversy surrounding PAC analysis, methodological steps were outlined in detail and justified by existing empirical research. Furthermore, follow-up analyses were conducted to establish the reliability of our results and to assess whether patterns of alpha-gamma PAC were driven by non-sinusoidal oscillations or insufficient data.

## 3. Methods

### 3.1 Participants

Data were collected from 16 participants (6 male, 10 female, mean age = 28.25, SD = 6.23). All participants had normal or corrected to normal vision and no history of neurological or psychiatric illness.

### 3.2 Experimental Procedures

All experimental procedures complied with the Declaration of Helsinki and were approved by the Aston University, Department of Life & Health Sciences ethics committee.

### 3.3 Paradigm

Participants performed an engaging sensory paradigm (figure 1), designed to elicit patterns of high-frequency oscillatory activity. Each trial started with a fixation period of 1500-3500ms, followed by the presentation of a visual grating or auditory binaural click train cue; however only visual data will be analysed in this article. The visual grating had a spatial frequency of 2 cycles/degree and was presented for 1500ms. Following this, participants were instructed to respond to the appearance of an alien stimulus (go) or astronaut stimulus (no-go) using a response pad. The astronaut/alien stimulus was presented for 500ms followed by a maximum response period of 1000ms. The accuracy of the response was conveyed through audio-visual feedback, followed by a 500ms baseline period. In total, the MEG recording lasted 12-13 minutes and included 64 trials with visual grating stimuli. Prior to MEG acquisition, the nature of the task was fully explained to participants and several practice trials were performed. Accuracy rates were above 95% for all participants indicating that the task was engaging and successfully understood.

**Figure 1:**
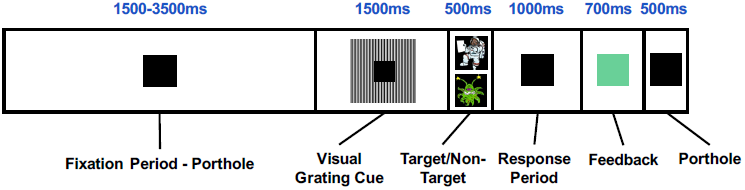
The structure of the engaging sensory paradigm, in which participants were instructed to respond after the appearance of the alien stimulus. Each trial started with a 1500-3500ms baseline period in which a square black box (the “porthole”) was centrally presented. This was followed by presentation of the visual grating (2 cycles/degree) around the central porthole for 1500ms. The target stimulus (alien) or non-target stimulus (astronaut) was then shown within the porthole for 500ms followed by a 1000ms response period. Correct or incorrect responses were conveyed to the participant through audio-visual feedback in which the porthole turned green (correct) or red (incorrect) and a correct/incorrect tone was played.

### 3.4 MEG Acquisition

MEG data were acquired using a 306-channel Neuromag MEG scanner (Vectorview, Elekta, Finland) made up of 102 triplets of two orthogonal planar gradiometers and one magnetometer. All recordings were performed inside a magnetically shielded room at a sampling rate of 1000Hz. Five head position indicator (HPI) coils were applied for continuous head position tracking, and visualised post-acquisition using an in-house Mat lab script. For MEG-MRI coregistration purposes three fiducial points, the locations of the HPI coils and 300-500 points from the head surface were acquired using the integrated Polhemus Fastrak digitizer.

Visual stimuli were presented on a screen located 86cm from participants (resulting in 2 cycles/degree for the visual grating), and auditory feedback through MEG-compatible headphones.

### 3.5 Structural MRI

A structural T1 brain scan was acquired for source reconstruction using a Siemens MAGNETOM Trio 3T scanner with a 32-channel head coil (TE=2.18ms, TR=2300ms, TI=1100ms, flip angle=9, 192 or 208 slices depending on head size, voxel-size = 0.8×0.8×0.8cm).

### 3.6 MEG MRI Coregistration and 3D Cortical Mesh Construction

MEG data were co-registered with participants MRI structural scan by matching the digitised head shape data with surface data from the structural scan (Jenkinson & Smith, 2001). The aligned MRI-MEG image was used to create a forward model based on a single-shell description of the inner surface of the skull (Nolte, 2003), using the segmentation function in SPM8 (Litvak et al., 2011). The cortical mantle was then extracted to create a 3D cortical mesh, using Freesurfer v5.3 (Fischl, 2012), and registered to a standard f_s__LR mesh, based on the Conte69 brain (Van Essen 2012), using an interpolation algorithm from the Human Connectome Project (Van Essen et al., 2012; instructions here: https://goo.gl/3HYA3L). Finally, the mesh was downsampled to 4002 vertices per hemisphere. Due to the extensive computation time involved in these procedures, all participant-specific cortical meshes are available to download in the /anat directory of the Figshare repository (see later).

### 3.7 Pre-processing

MEG data were pre-processed using Maxfilter (temporal signal space separation, .9 correlation), which supresses external sources of noise from outside the head (Taulu & Simola, 2006).

Further pre-processing steps were performed in Matlab 2014b using the open-source Fieldtrip toolbox v20161024 (Oostenveld et al., 2010; script: 1_preprocessing_elektra_frontiers_PAC.m). Firstly, for each participant the entire recording was band-pass filtered between 0.5-250Hz (Butterworth filter, low-pass order 4, high-pass order 3) and band-stop filtered (49.5-50.5Hz; 99.5-100.5Hz) to remove residual 50Hz power-line contamination and its harmonics. Data were then epoched into segments of 4000ms (2000ms pre, 2000ms post stimulus onset) and each trial was demeaned and detrended. Trials containing artefacts (SQUID jumps, eye-blinks, head movement, muscle) were removed if the trial-by-channel (magnetomer) variance exceeded 8x10^−23^, resulting in an average of 63.5 trials per condition, per participant. Site-specific MEG channels containing large amounts of non-physiological noise were removed from all analyses (MEG channels: 0111, 0332, 2542, 0532).

### 3.8 Source Analysis

Source analysis was conducted using a linearly constrained minimum variance beamformer (LCMV) (Van Veen, van Drongelen, Yuchtman, & Suzuki, 1997), which applies a spatial filter to the MEG data at each vertex of the 3D cortical mesh, in order to maximise signal from that location whilst attenuating signals elsewhere. Beamforming weights were calculated by combining the covariance matrix of the sensor data with leadfield information. Due to rank reduction following data cleaning with Maxfilter, the covariance matrix was kept at a rank which explained 99% of the variance. For all analyses, a common filter was used across baseline and grating periods, and a regularisation parameter of lambda 5% was applied.

Due to prior interest in the gamma and alpha-bands (Hoogenboom, Schoffelen, Oostenveld, Parkes, & Fries, 2006; Michalareas et al., 2016; Muthukumaraswamy, Singh, Swettenham, & Jones, 2010), the visual data were band-pass filtered (Butterworth filter) between 40-60Hz (gamma) and 8-13Hz (alpha), and source analysis was performed separately for each frequency band. To capture induced rather than evoked visual activity, a period of 300-1500ms following stimulus onset was compared with a 1200ms baseline period. The change in oscillatory power for each vertex was averaged across participants, interpolated onto a 3D mesh provided by the Human Connectome Project (Van Essen, 2012), and thresholded at a value which allowed the prominent patterns of power changes to be determined.

### 3.9 Extracting Area V1 Time-series

Trial time-courses were extracted from bilateral visual area V1, defined using a multi-modal parcellation from the Human Connectome Project, which combined retinotopic mapping, T1/T2 structural MRI and diffusion-weighted MRI to accurately define the boundaries between cortical areas (Glasser et al., 2016; Figure 3C). The downsampled version of this atlas can be found in the parent directory of the Figshare repository (see later). To obtain a single spatial filter from this region, we performed a principle components analysis (PCA) on the concatenated filters from 182 vertices of bilateral V1, multiplied by the sensor-level covariance matrix, and extracted the first component. The sensor-level data was then multiplied by this spatial filter to obtain a V1-specific “virtual electrode”, and the change in oscillatory power between grating and baseline periods was calculated from 1-100Hz, using a sliding window of 500ms and fixed frequency smoothing (±8Hz) (Hoogenboom, Schoffelen, Oostenveld, Parkes, & Fries, 2006). It is important to note that while we decided to use a multimodal atlas, visual area V1 virtual electrode time-series could also be defined using a more standard volumetric approach, for example the AAL atlas, which is included in the Fieldtrip toolbox (Oostenveld et al., 2010).

### 3.10 Phase Amplitude Coupling (PAC) Analysis

V1 time-courses were examined for changes in alpha-gamma phase amplitude coupling (PAC). The general procedure is outlined in Figure 2. The first step was to obtain estimates of low frequency phase (*f*_p_) and high frequency amplitude (*f*_a_) for each trial using a fourth order, two-pass Butterworth filter, and then applying the Hilbert transform (Le Van Quyen et al., 2001). To avoid sharp edge artefacts, which can result in spurious PAC (Kramer, Tort, & Kopell, 2008), the first 800ms and last 500ms of each trial was discarded.

**Figure 2:**
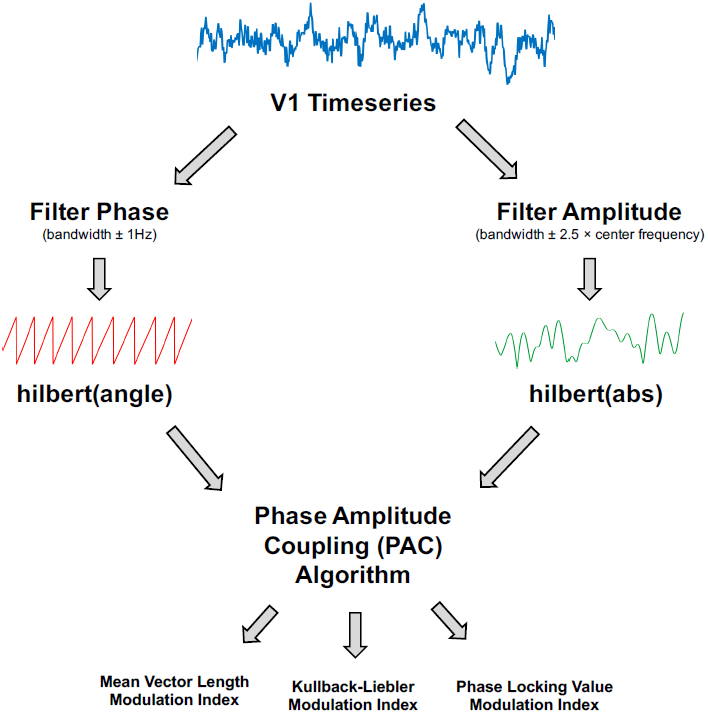
Illustration of the phase amplitude coupling (PAC) analysis procedure. The V1 time-series were filtered to obtain estimates of phase and amplitude, using a narrow (±1Hz) bandwidth for the phase and a variable bandwidth (±2.5 times the centre frequency) for the amplitude. Phase and amplitude information were obtained via the Hilbert transform. The coupling between phase and amplitude was then quantified using Mean Vector Length, Kullback-Leiber or Phase Locking Value algorithms to produce a Modulation Index value.

The bandwidth of the filter used to obtain *f*_p_ and *f*_a_ is a crucial parameter in calculating PAC (Aru et al., 2015). The filters for extracting *f*_a_ need to be wide enough to capture the centre frequency ± the modulating *f*_p_. So, for example, to detect PAC between *f*_p_ = 13Hz and *f*_a_ = 60Hz, requires a *f*_a_ bandwidth of at least 13Hz [47 73]. If this condition is not met, then PAC cannot be detected even if present (Dvorak & Fenton, 2014). We therefore decided to use a variable bandwidth, defined as ±2.5 times the center frequency (e.g. for an amplitude of 60Hz, the bandwidth was 24Hz either side [36 84]), which has been shown to improve the ability to detect PAC (Berman et al., 2012; Voloh et al., 2015). For alpha-band phase (maximum 13Hz), this allowed us to calculate PAC for amplitudes above 34Hz. The bandwidth for f_p_ was kept narrow (1Hz ± the center frequency), in order to extract sinusoidal waveforms. Furthermore, each trial was visually inspected to confirm that the *f*_p_ filtered oscillations were sinusoidal in nature.

Next, the coupling between *f*_p_ and *f*_a_ was quantified using four common PAC approaches^1^: the Mean-Vector Length modulation index, originally described in Canolty et al., (2006); the Mean-Vector Length modulation index described in Özkurt & Schnitzler (2011); the phase-locking value modulation index described in Cohen (2008); and the Kullback-Lieber modulation index described in Tort, Komorowski, Eichenbaum, & Kopell (2010a). These approaches were selected due to their popularity in the MEG/EEG PAC literature (e.g. Bonnefond & Jensen, 2015; Cho et al., 2015; Khan et al., 2013; Mathewson et al., 2011), and to demonstrate the diversity of PAC results based on the algorithm selected.

The mean vector length modulation index (MVL-MI-Canolty) approach estimates PAC by combining phase (ϕ) and amplitude information to create a complex-valued signal: f_*a*_*e*^*i*(*ϕf*_*p*_)^(Canolty et al., 2006), in which each vector corresponds to a certain time-point (N). If the resulting probability distribution function is non-uniform, this suggests a coupling between *f*_p_ and *f*_a_, which can be quantified by taking the length of the average vector.

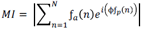

However, MI values from the MVL-MI-Canolty algorithm have been shown to partly reflect the power of *f*_a_ oscillations, rather than their coupling (Canolty & Knight, 2010). Therefore, as an alternative to surrogate data, we applied a MVL-MI algorithm from Özkurt & Schnitzler (2011), which includes a normalisation factor corresponding to the power of *f*_a_. Özkurt & Schnitzler (2011) suggest that their algorithm is more resilient to measurement noise, and is therefore highly relevant for MEG data, which has an inherently lower signal-to-noise ratio compared with invasive electrophysiological recordings (Goldenholz et al., 2009).

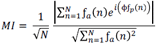

The PLV-MI-Cohen approach assumes that if PAC is present, the envelope of *f*_a_ should oscillate at the frequency corresponding to *f*_p_. The phase of *f*_a_ envelope can be obtained by applying the Hilbert transform (angle): ϕ*f*_a_. The coupling between the low-frequency ϕf_p_ phase values and the phase of the amplitude envelope, ϕf_a_, can be quantified by calculating a phase locking value (PLV), in much the same way as determining phase synchronisation between electrophysiological signals.

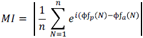

Finally, the KI-MI-Tort approach estimates PAC by quantifying the amount of deviation in amplitude-phase distributions. This involves breaking *f*_p_ into 18 bins, and calculating the mean amplitude within each phase bin, normalised by the average value across all bins. The modulation index is calculated by comparing the amplitude-phase distribution (P) against the null hypothesis of a uniformly amplitude-phase distribution (Q).

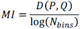

Mathematically, this is computed using the Kullbeck-Leiber distance (D), related to Shannon’s entropy.

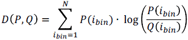

Using these four approaches (MVL-MI-Canolty; MVL-MI-Özkurt; KL-MI-Tort; PLV-MI-Cohen) we calculated PAC between phases 7-13Hz (in 1 Hz steps) and amplitudes 34-100Hz (in 2Hz steps), for the time-period 300-1500ms following grating presentation and a 1200ms baseline period. PAC values were calculated separately for each trial and then averaged to obtain a single MI value per amplitude and phase. This was repeated using surrogate data, created by shuffling trial and phase-carrying information (200 surrogates), to normalise MI values. On a PC with 32GB of RAM, and Intel(R) Core™ i7-4790 processor, the computation time for these procedures was 4.5 hours.

To assess changes in the strength of PAC between the grating and baseline periods, the comodulograms were compared using non-parametric cluster-based statistics, which have been shown to adequately control the type-I error rate for electrophysiological data (Maris & Oostenveld, 2007). First, an uncorrected dependent-samples t-test was performed (grating versus baseline), and all MI values exceeding a 5% significance threshold were grouped into clusters. The maximum t-value within each cluster was carried forward. Next, a null distribution was obtained by randomising the condition label (grating/baseline) 1000 times and calculating the largest cluster-level t-value for each permutation. The maximum t-value within each original cluster was then compared against this null distribution, with values exceeding a threshold of p<.05 deemed significant.

### 3.11 Sinusoidal Oscillations

One major issue in cross-frequency coupling analysis is the presence of non-sinusoidal sawtooth-like oscillations (Cole et al., 2017; Jensen et al., 2016), which can result in spurious estimates of PAC (Lozano-Soldevilla et al., 2016). This property of oscillations can be quantified by calculating the time taken from trough to peak (rise-time), peak to trough (decay-time), and the ratio between these values (Cole & Voytek, 2017; Dvorak & Fenton, 2014). We therefore calculated this ratio for the visual V1 data from 7-13Hz, to check for differences in non-sinusoidal oscillations between grating and baseline periods.

### 3.12 Simulated PAC Analysis

To investigate the validity of the four PAC approaches, we constructed 1.2 seconds of simulated data with known alpha-gamma PAC (f_p_ = 10Hz; *f*_a_ = 50-70Hz; code adapted from Kramer et al., (2008) and Özkurt & Schnitzler (2011)) and added a random level of noise (signal-to-noise ratio > -11.5Db). Comodulograms were produced using the four PAC algorithms on 64 trials of simulated data. Using the same code, we also investigated how the four algorithms were affected by trial length (0.1-10s in 0.1 second steps).

### 3.13 Analysis Code & Data Sharing

MEG data are available to download online at Figshare (https://doi.org/10.6084/m9.figshare.c.3819106.v1), along with participant-specific 3D cortical meshes. Access to the raw structural MRI data will be granted upon reasonable request and ethical approval from Aston University Life & Health Sciences ethics committee. MATLAB code used for all analyses has been made available in the supplementary materials and online (https://github.com/neurofractal/sensory_PAC), including the four PAC algorithms, which can be applied to electrophysiological data arranged in the standard Fieldtrip format (Oostenveld et al., 2010). Successful use of the scripts requires the user to have at least a basic understanding of MATLAB, signal processing, and the methodological complexities surrounding PAC. We therefore direct the reader to a number of excellent reviews and empirical papers (Aru et al., 2015; Canolty et al., 2006; Canolty & Knight, 2010; Hyafil et al., 2015; Jensen & Colgin, 2007).

## 4. Results

### 4.1 Source Localisation

In order to establish patterns of oscillatory power changes following presentation of the visual grating, gamma-band (40-60Hz) and alpha-band power (8-13Hz) were localised for a 300-1500ms period post-stimulus presentation. Results for the gamma-band (figure 3A), show an increase in oscillatory power which localises to the ventral occipital cortex (Hoogenboom et al., 2006). Results for the alpha band (figure 3B) showed a general decrease in power, located primarily in occipital areas, but extending into temporal and parietal regions. The more widespread spatial pattern could reflect on-going upstream processes triggered by the appearance of the grating, for example anticipation of the upcoming target (Stenner, Bauer, Haggard, Heinze, & Dolan, 2014).

**Figure 3:**
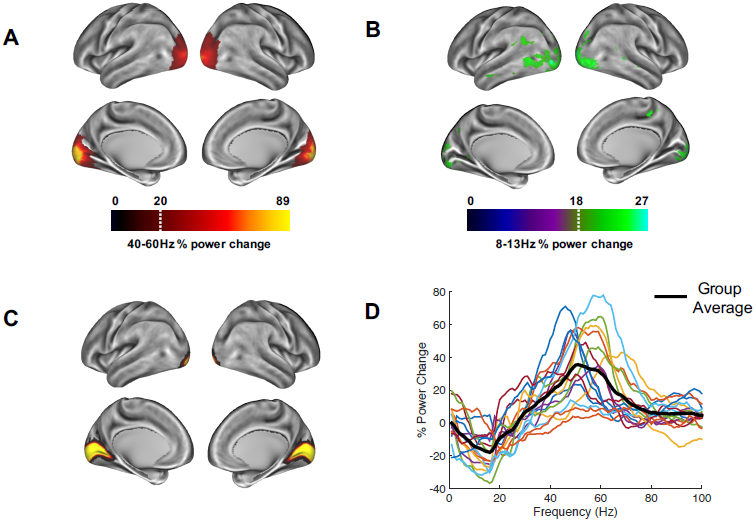
Whole-brain oscillatory power changes following the presentation of the visual grating are marked by **(A)** increases in the gamma-band (40-60Hz) and **(B)** decreases in the alpha-band (8-13Hz), localised primarily in the ventral occipital cortex (script: 2_get_source_power.m). Power maps were thresholded at a value which allowed prominent patterns of power changes to be determined, indicated by the white dotted line. Time-courses were extracted from bilateral visual area V1 (script: 3_get_VE_frontiers_PAC.m), defined using the atlas region shown in **(C)** from the HCP-MMP 1.0 parcellation (Glasser et al., 2016). **(D)** These V1 responses showed reductions in alpha/beta power and increases in gamma-band (40-70Hz) power (script: 4_calc_pow_change.m).

### 4.2 Visual Area V1 Power Changes

Time courses from area V1 were extracted (figure 3C), and the change in oscillatory power between grating and baseline periods from 1-100Hz was calculated (figure 3D). Whilst results show individual variability in peak frequencies and the strength of oscillatory power, on average, activity within visual area V1 displays a reduction in alpha/beta power (8-20Hz), and an increase in gamma power (40-70Hz). The MEG data, therefore display well-established patterns of alpha and gamma-band event-related synchronisation and desynchronisation within visual area V1 (Bonnefond & Jensen, 2015; Hoogenboom et al., 2006; Michalareas et al., 2016), which is a crucial first step in calculating reliable estimates of PAC (Aru et al., 2015).

### 4.3 Alpha-Gamma PAC

Visual area V1 responses were next examined for changes in alpha-gamma PAC. Specifically, we set out to test whether the coupling between alpha-band phase and gamma-band amplitude was altered during presentation of the visual grating. Phase-amplitude comodulograms were produced between a range of phase frequencies (7-13Hz) and amplitude frequencies (34-100Hz), using the four algorithms described in Methods: MVL-MI-Canolty; MVL-MI-Özkurt; PLV-MI-Cohen and KL-MI-Tort. Grating and baseline comodulograms were compared using cluster-based non-parametric statistics (Maris & Oostenveld, 2007).

Results are shown in Figure 4A. Using the MVL-MI-Canolty algorithm, there was a significant increase in alpha-gamma PAC over a large proportion of the comodulogram, between 40-100Hz and 7-13Hz, with a peak at 50-70Hz amplitude and 9-10Hz phase. This large area of significantly increased PAC is likely to reflect, in part, power increases in the gamma-band (Canolty et al., 2006). The alternative MVL-MI-Özkurt algorithm, which normalises MI values by the high-frequency oscillatory power, displayed a smaller area of significant coupling, with increased PAC between an amplitude of 50-70Hz and phase of 10Hz. There was also a similar cluster of significantly increased PAC between 9-11Hz and 50-70Hz using the PLV-MI-Cohen approach. The KL-MI-Tort results showed clusters of increased PAC between amplitudes of 50-100Hz and phases of 9-10Hz, but decreased PAC between amplitudes of 60-90Hz and phases of 12-13Hz. However, none of these clusters passed a significance threshold of p<0.05 (two-tailed). Similar results were obtained after normalising MI values with surrogate data (Figure 4B).

**Figure 4:**
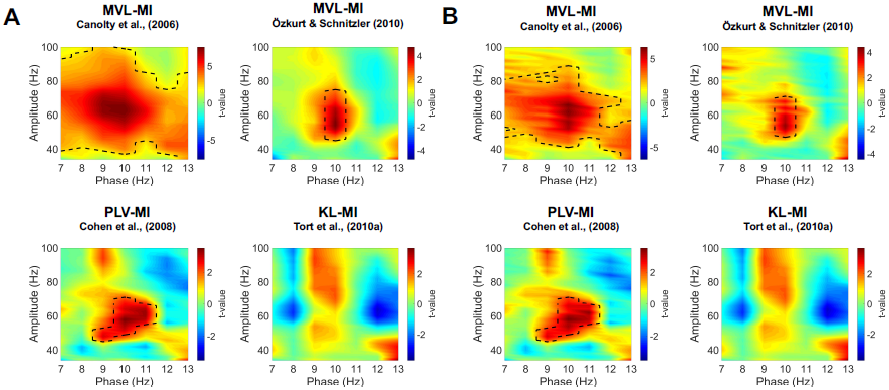
Phase-amplitude comodulograms produced by statistically comparing modulation index (MI) values from 300ms-1500ms post-grating onset to a 1200ms baseline period, using four separate approaches (script: 5_visual_PAC_four_methods.m). Comodulograms for (A) raw MI values and (B) MI values normalised by surrogate data are shown separately. The black dotted line represents significantly different phase-amplitude coupling frequencies (p<.05; for details of non-parametric cluster-based statistics see Methods).

### 4.4 Non-Sinusoidal Oscillations

To determine whether our alpha-gamma PAC results were driven by differences in the sinusoidal properties of oscillations between baseline and grating periods, the ratio between oscillatory rise-time and decay-time was calculated (6_check_non_sinusoidal.m). For the alpha phase frequencies (7-13Hz), there was no difference in this ratio (all frequencies p>.05), suggesting that our results are unlikely to be caused by increased non-sinusoidal sawtooth-like properties of alpha oscillations during stimulus period compared to baseline.

### 4.5 Simulated PAC

To further validate our PAC results, we generated simulated data with known alpha-gamma coupling (10-11Hz phase, 50-70Hz amplitude). Using the same MATLAB code as for the MEG data, we were able to successfully detect this alpha-gamma PAC using the MVL-MI-Canolty, MVL-MI-Özkurt, PLV-MI-Cohen and KL-MI-Tort algorithms (Figure 5A). By varying the trial length of the simulated data, we found that PAC values were affected by trial length, with data segments under 1 second producing artificially inflated PAC (Figure 5B).

**Figure 5:**
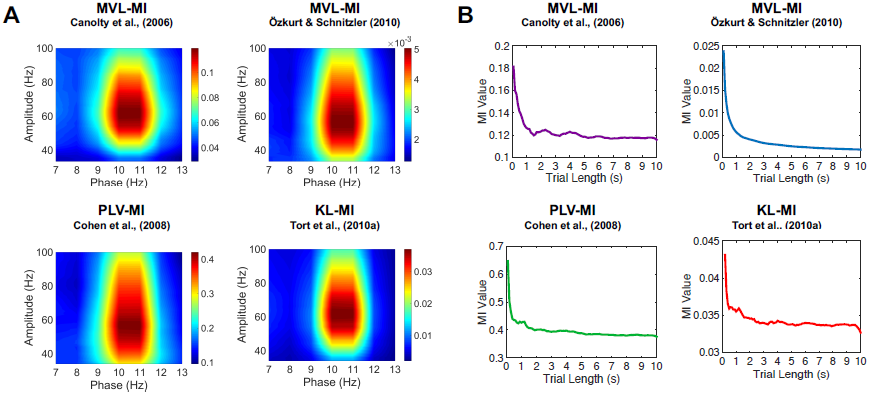
Results of the simulated PAC analysis. **(A)** Phase-amplitude comodulograms produced using the MVL-MI-Canolty, MVL-MI-Özkurt, PLV-MI-Cohen and KL-MI-Tort algorithms were able to successfully detect the simulated coupling between 10Hz phase and 50-70Hz amplitude. **(B)** The coupling between 10Hz phase and 60Hz amplitude was calculated as a function of simulated data trial length. For trial data under 1 second, all four algorithms produced artificially inflated PAC (script: 7_simulated_PAC_analysis.m).

## 5. Discussion

This article has outlined various steps involved in the detection and validation of phase amplitude coupling (PAC) in a visual MEG dataset (data shared at: https://doi.org/10.6084/m9.figshare.c.3819106.v1), utilising the open-source Fieldtrip toolbox (Oostenveld et al., 2010) and customised Matlab scripts (all scripts shared at: https://github.com/neurofractal/sensory_PAC). We first confirmed that presentation of the visual grating was accompanied by decreases in alpha power (8-13Hz) and increases in gamma power (>40Hz) within visual area V1. Although this may seem redundant given the wealth of evidence for alpha and gamma oscillations in visual processing (Bonnefond & Jensen, 2015; Hoogenboom et al., 2006; Michalareas et al., 2016), it is crucial to establish clear increases/decreases in the power spectrum at two distinct frequencies as a first step in MEG-PAC analysis (Aru et al., 2015; Hyafil et al., 2015). Using four PAC algorithms, we showed that visual responses obtained from area V1 displayed a general increase in alpha-gamma PAC as expected (Bonnefond & Jensen, 2015; Spaak et al., 2012; Voytek et al., 2010). However, it is important to note that specific patterns of coupling depended on the algorithm selected. The MVL-MI-Canolty algorithm showed large increases in PAC during the grating period, covering almost the entire alpha & gamma frequency ranges, most likely as a result of MI values being biased by increases in high-frequency power following presentation of the visual grating (Canolty et al., 2006). This approach is therefore less suitable for detecting PAC between separate periods of data and/or trials. The MVL-MI-Özkurt algorithm, which normalises the MI value by high amplitude power, along with the PLV-MI-Cohen algorithm produced a much more constrained pattern of significant alpha-gamma PAC, with peaks between 9-11Hz phase and 50-70Hz amplitude. Whilst the KL-MI-Tort approach also showed a general increase in alpha-gamma PAC around 9-11Hz, none of the phase-amplitude clusters reached significance. This may be due to the relatively short number of trials used in the experiment, the low signal-to-noise ratio of MEG recordings (Goldenholz et al., 2009), variations in the peak alpha and gamma oscillatory frequencies (Muthukumaraswamy, Edden, Jones, Swettenham, & Singh, 2009), combined with the fact that the KL-MI-Tort approach is relatively conservative (van Driel, Cox, & Cohen, 2015). More generally, it is important to emphasise that all four PAC metrics are highly sensitive to a range of factors (Aru et al., 2015; Dvorak & Fenton, 2014), which are often hard to control (Berman et al., 2012), resulting in both type I and type II statistical errors.

One such issue is the presence of non-sinusoidal sawtooth-like oscillations in electrophysiological data, which can result in spurious PAC (Lozano-Soldevilla et al., 2016), especially when phase is obtained with wide band-pass filters. By computing the ratio between rise-time and decay-time of alpha oscillations within area V1, we showed that non-sinusoidal oscillations did not differ between baseline and grating periods, and are unlikely to account for our results. Another issue in trial-based PAC analysis is data length, with some previous reports suggesting that 10 seconds or more is required for detecting theta-gamma coupling (Aru et al., 2015; Dvorak & Fenton, 2014). However, using simulated alpha-gamma PAC we determined that 1 second of data was sufficient to obtain stable estimates. We encourage the reader to run similar follow-up analyses after finding significant PAC to check for spurious coupling caused by, for example, non-sinusoidal oscillations (Jensen et al., 2016; Lozano-Soldevilla et al., 2016) and/or insufficiently long trials (Dvorak & Fenton, 2014).

### 5.1 Practical Considerations for PAC analysis

Cross-frequency coupling is gaining significant interest within the electrophysiological community (Aru et al., 2015; Canolty & Knight, 2010; Dvorak & Fenton, 2014; Hyafil et al., 2015), and therefore it is important for researchers to consider the methodological pitfalls and caveats which commonly arise during PAC analysis. Firstly, due to the presence of edge artefacts at the start and end of time-series created by bandpass filtering, which can result in artefactual PAC (Kramer et al., 2008), sufficient padding should be included around trials. Concatenating data from separate trials to create longer data segments results in similar edge artefacts (Kramer et al., 2008), and should be avoided. Secondly, if the bandwidth of the filter used to extract the amplitude does not contain the side-bands of the modulating phase frequency, PAC cannot be detected even if present (Dvorak & Fenton, 2014). The use of a variable band-pass filter which scales with amplitude, can alleviate this issue and improve the sensitivity of detecting PAC (Berman et al., 2012; Voloh et al., 2015). Thirdly, periods which contain non-stationary periods should be avoided. This includes sensory evoked potentials which induce correlations between frequency bands via phase reset (Sauseng et al., 2007), and can be misinterpreted as PAC (Aru et al., 2015). For this reason, we did not analyse the first 300ms following visual grating presentation, due to the presence of visual evoked potentials (Di Russo, Martínez, Sereno, Pitzalis, & Hillyard, 2002). Fourth, given that PAC algorithms produce values ranging from 0 to 1, data are commonly not normally distributed, and therefore the use of non-parametric statistics is paramount. Whilst surrogate data are often employed (Aru et al., 2015; Tort et al., 2010a), this may not be possible where data are organised into short trials and temporal correlations between surrogate and true time-series are high (Dvorak & Fenton, 2014). Therefore, to assess changes in PAC, using a baseline period or contrasting between conditions, combined with non-parametric statistics may prove to be a useful alternative for sensory neurocognitive research.

### 5.2 Limitations

This study has compared four PAC algorithms (Canolty et al., 2006; Cohen, 2008; Özkurt & Schnitzler, 2011; Tort et al., 2010a), which are among the most commonly used approaches in sensory EEG/MEG research (Bonnefond & Jensen, 2015; Cho et al., 2015; Khan et al., 2013; Mathewson et al., 2011). However these only comprise a small subset of the available algorithms designed to quantify PAC (Canolty & Knight, 2010; Hyafil et al., 2015). There have also been advances in measuring transient changes in PAC (Dvorak & Fenton, 2014), directed PAC (Jiang, Bahramisharif, van Gerven, & Jensen, 2015) and algorithms designed for spontaneous neural activity (Florin & Baillet, 2015; Weaver et al., 2016). A more comprehensive evaluation of algorithms and their application to real-world electrophysiological data is beyond the scope of this article, but would nevertheless benefit the field of cross-frequency coupling. Secondly, in order to detect alpha-gamma PAC within visual area V1, we used a broad filter bandwidth, defined as ±2.5 times the amplitude centre-frequency. Consequently, the alpha-gamma comodulograms will be unable to differentiate between adjacent gamma sub-bands, which have been proposed to fulfil differing neurocognitive roles (Bosman, Lansink, & Pennartz, 2014; Buzsáki & Wang, 2012), and patterns of PAC (Vaz et al., 2017). However, for the visual MEG data presented here, there was only an increase in gamma power within one band (40-70Hz), and therefore the smearing of adjacent sub-bands is unlikely. Finally, we have focussed on PAC within the visual cortex, which is known to display highly sinusoidal alpha oscillations (Tort et al., 2010b). However, there are many examples of non-sinusoidal brain oscillations caused by physiological neuronal spiking patterns (Fontanini & Katz, 2005), including hippocampal theta (4-8Hz) and sensorimotor mu (9-11Hz) rhythms (Lozano-Soldevilla et al., 2016; Scheffer-Teixeira & Tort, 2016), which are indicative of behaviour and disease states (Cole & Voytek, 2017). Therefore, whilst non-sinusoidal oscillations generate spurious PAC, this does not mean that these oscillations are uninteresting, but simply that common PAC algorithms, such as the ones employed in this article, are ill-suited for these scenarios. Where non-sinusoidal oscillations are present, PAC analysis could proceed by correcting for non-uniform phase distributions (e.g van Driel, Cox, & Cohen, 2015) in order to disentangle nested oscillations from neural spiking (Vaz et al., 2017).

### 5.3 Conclusion

In conclusion, we have outlined the key analysis steps for detecting changes in alpha-gamma PAC during sensory processing, using an example visual MEG dataset. While alpha-gamma PAC was shown to increase, the specific patterns of alpha-gamma coupling depended upon the specific algorithm employed. Follow-up analyses showed that these results were not driven by non-sinusoidal oscillations or insufficient data. In future, we hope that a variety of PAC algorithms will be implemented alongside existing open-source MEG toolboxes (Gramfort et al., 2014; Oostenveld et al., 2010; Tadel, Baillet, Mosher, Pantazis, & Leahy, 2011), with detailed guidance and advice, so that PAC can form a natural analysis step in electrophysiological research.

## 6. Author Contributions

RS, KK & GR co-designed the study and wrote the manuscript. RS collected the data, carried out the analysis and organised the data and code for sharing.

## 8. Acknowledgments

We wish to thank: the volunteers who gave their time to participate in this study; Jan-Mathijs Schoffelen for providing MATLAB code; Gerard Gooding-Williams and Shu Yau for help with MRI data acquisition; Diandra Brkic for helpful comments on the manuscript and the Wellcome and Dr Hadwen Trusts for supporting MEG scanning costs.

## Footnotes

1 Due to inconsistent naming practices, we refer to the quantitative value of PAC as the modulation index (MI) across all four approaches.

